# Visible light-induced specific protein reaction delineates early stages of cell adhesion

**DOI:** 10.1101/2023.07.21.549850

**Authors:** Rolle Rahikainen, Susan K. Vester, Paula Turkki, Chasity P. Janosko, Alexander Deiters, Vesa P. Hytönen, Mark Howarth

## Abstract

Light is well established for control of bond breakage, but not for control of specific bond formation in complex environments. We previously engineered diffusion-limited reactivity of SpyTag003 peptide with its protein partner SpyCatcher003 through spontaneous transamidation. This system enables precise and irreversible assembly of biological building blocks, with applications from biomaterials to vaccines. Here, we establish a system for rapid control of this amide bond formation with visible light. We have generated a caged SpyCatcher003, which allows light triggering of covalent bond formation to SpyTag003 in mammalian cells. Photocaging is achieved through site-specific incorporation of an unnatural coumarin-lysine at the reactive site of SpyCatcher003. We showed uniform specific reaction in cell lysate upon light activation. We then used the spatiotemporal precision of a 405 nm confocal laser for uncaging in seconds, probing the earliest events in mechanotransduction by talin, the key force sensor between the cytoskeleton and extracellular matrix. Reconstituting talin induced rapid biphasic extension of lamellipodia, revealing the kinetics of talin-regulated cell spreading and polarization. Thereafter we determined the hierarchy of recruitment of key components for cell adhesion. Precise control over site-specific protein reaction with visible light creates diverse opportunities for cell biology and nanoassembly.

## MAIN TEXT

Living systems display exquisite precision in their organization and rapid adaptation. Chemical biology aims to exert control over cell or organism behavior, but most methods act over hours to days (genetic modification) or lack spatial control (pharmacological manipulation).^1,2^ However, light allows rapid and precise subcellular responses, e.g. optogenetics to modulate membrane gradients for electrical signaling.^3^ In the area of protein interactions, interactions can be switched by visible light, using phytochrome or light-oxygen voltage (LOV) domains.^1^ We have endeavored to develop protein-protein interactions that extend beyond typical stability, through genetically-encoded irreversible ligation.^4^ SpyTag003 is a peptide that we have engineered for rapid transamidation with its protein partner SpyCatcher003 (Fig. 1A).^5^ Reaction proceeds close to the diffusion limit, occurs under diverse conditions, and is efficient in numerous cellular systems.^5,6^ Tag/Catcher bioconjugation has been employed in biomaterials, vaccine assembly, and antibody functionalization.^4,7,8^ SpyTag003/SpyCatcher003 has also been useful inside cells, including recruitment of epigenetic modifiers or enzyme channeling.^4,9,10^ Previously, an engineered LOV domain allowed photocontrol of SpyTag/SpyCatcher, although there was gradual transamidation even in the dark-state.^11^ To enable highly switchable covalent reaction, here we employ site-specific incorporation of unnatural amino acids.^12^ Photoreactive amino acids like benzoylphenylalanine trap complexes after UV activation,^13^ which is powerful to identify unknown complexes but not ideal for targeted bridging.^13^ Individual amino acids can also be photocaged^12,14,15^ and K31 is the key reactive residue on SpyCatcher003 (Fig. 1A).^5^ We focused our efforts on the unnatural amino acid 7-hydroxycoumarin lysine (HCK) (Fig. 1B) because uncaging in the visible spectrum (Fig. 1C) would reduce phototoxicity that is particularly serious in the UV range.^14,16,17,18^ Here, we establish caging of SpyCatcher003 using unnatural coumarin-lysine amino acid and its uncaging with 405 nm light for spatiotemporal control in living cells, to reveal early steps in mammalian cell adhesion.

**Figure 1.**
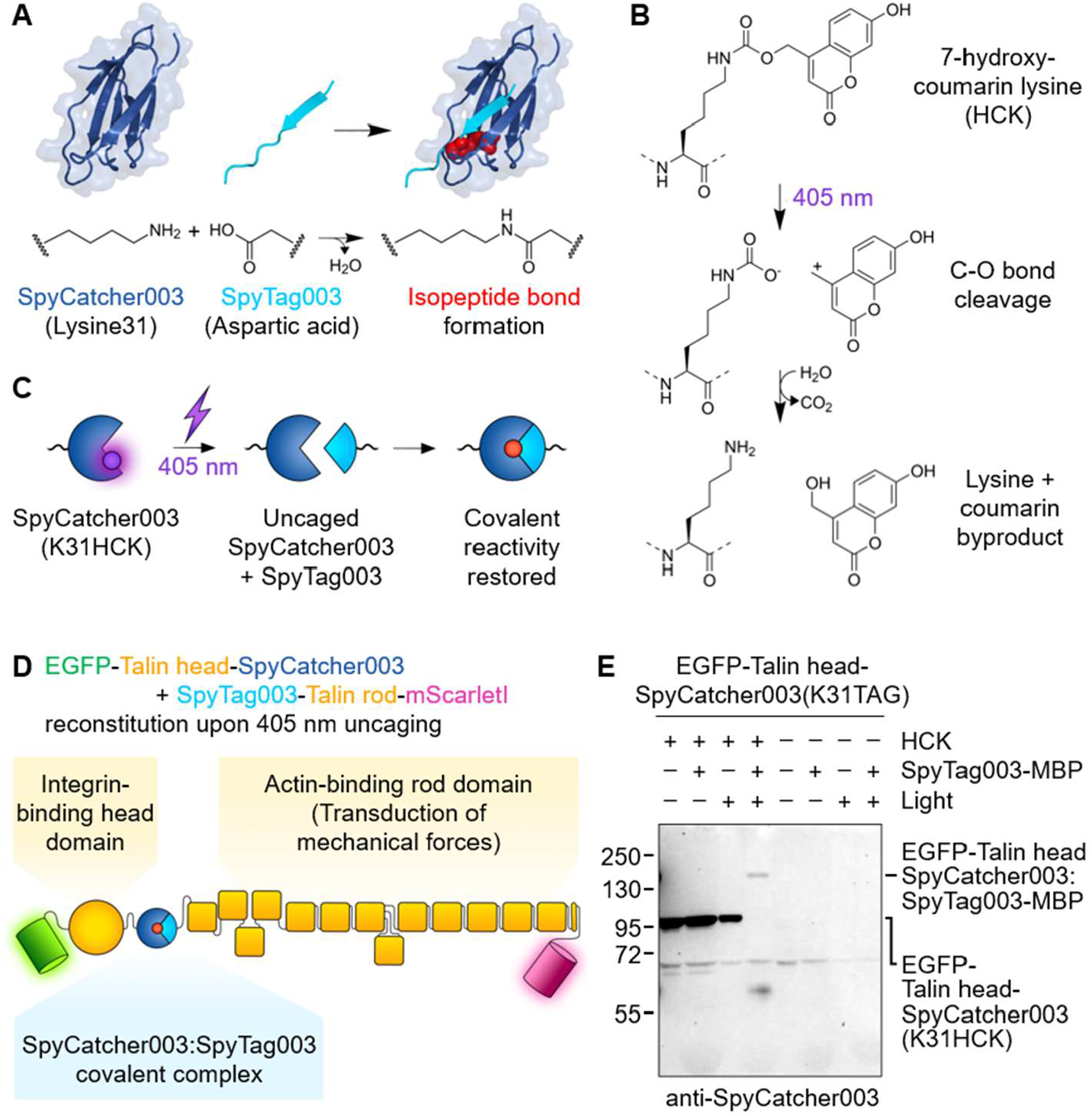
SpyCatcher003 photocaging with 7-hydroxycoumarin lysine. (A) Schematic of SpyTag003/SpyCatcher003 reaction. Lysine on SpyCatcher003 (dark blue) and aspartic acid on SpyTag003 (cyan) form a spontaneous isopeptide bond (reacted side-chains shown as red spheres), based on PDB 4MLI. (B) Schematic of light-induced cleavage. Dotted lines indicate the rest of the polypeptide. (C) Schematic for SpyCatcher003(K31HCK) uncaging. (D) Split-talin reconstitution using SpyCatcher003(K31HCK). (E) Covalent reactivity upon SpyCatcher003(K31HCK) photoactivation. Lysates of HEK293T cells transfected with EGFP-Talin head-SpyCatcher003(K31HCK) with or without HCK and light activation were mixed with SpyTag003-MBP, before Western blotting with anti-SpyCatcher003.

To establish our uncaging approach, we co-transfected the human cell-line HEK293T with HCK tRNAs and HCK tRNA synthetase (HCK RS)^16,19^ along with our protein of interest, to show that expression depended on the unnatural amino acid. Our initial construct contained the N-terminal region of transferrin receptor, SpyCatcher003 with an amber stop codon at K31 (K31TAG), and superfolder green fluorescent protein (sfGFP). Based on Western blotting, we optimized the dose of HCK and the ratio of the SpyCatcher003 construct to HCK RS (Fig. S1A). We then incorporated SpyCatcher003(K31TAG) in a split talin construct (Fig. 1D). We detected expression by blotting with antiserum to SpyCatcher003 (Fig. 1E) or anti-EGFP (Fig. S1B). We tested reactivity by adding SpyTag003-maltose binding protein (MBP) to cell lysate and monitoring bond formation by Western blot. HCK-caged SpyCatcher003 did not react with SpyTag003-MBP until lysate was treated with UV, consistent with effective photocaging of SpyCatcher003 (Fig. 1E, Fig. S1B). Without HCK, no SpyCatcher003 expression was detected, indicating that the stop codon led to chain termination (Fig. 1E). To understand practicality for selective uncaging, we assessed uncaging by ambient light. Room lighting or UK sunlight for 120 min did not lead to substantial uncaging in cell lysate in microcentrifuge tubes (Fig. S1C).

We applied photocontrolled reaction to gain insight into cell adhesion. Talin bridges the cytoplasmic domain of β-integrin to the actin cytoskeleton, completing the mechanical linkage between a cell and its surroundings.^20^ Talin functions as a molecular clutch that is required for cell spreading (Fig. 2A).^21–23^ Talin changes conformation in response to force, regulating association and release of multiple proteins involved in the cell’s response to mechanical cues.^24^ Talin recruitment has been previously controlled by an elegant strategy using rapamycin as a cell-permeable small molecule to reconstitute FRB- and FKBP-split talin fragments.^25^ However, this approach lacks subcellular spatial resolution and was only tested to withstand force of 4 pN,^25^ which may not resist sustained cytoskeletal tension.^26^ Split talin reconstitution using LOV domains would allow spatial control, but depends on continuous 488 nm illumination and has limited complex stability.^27^ Because of the complex structure and natural turnover of adhesion structures, estimating the impact of such non-covalently reconstituted talin on adhesion function is challenging. Rapid light-mediated induction of covalent talin reconstitution would allow precise control over early phases of adhesion formation, to decipher molecular details of talin-dependent processes.

**Figure 2.**
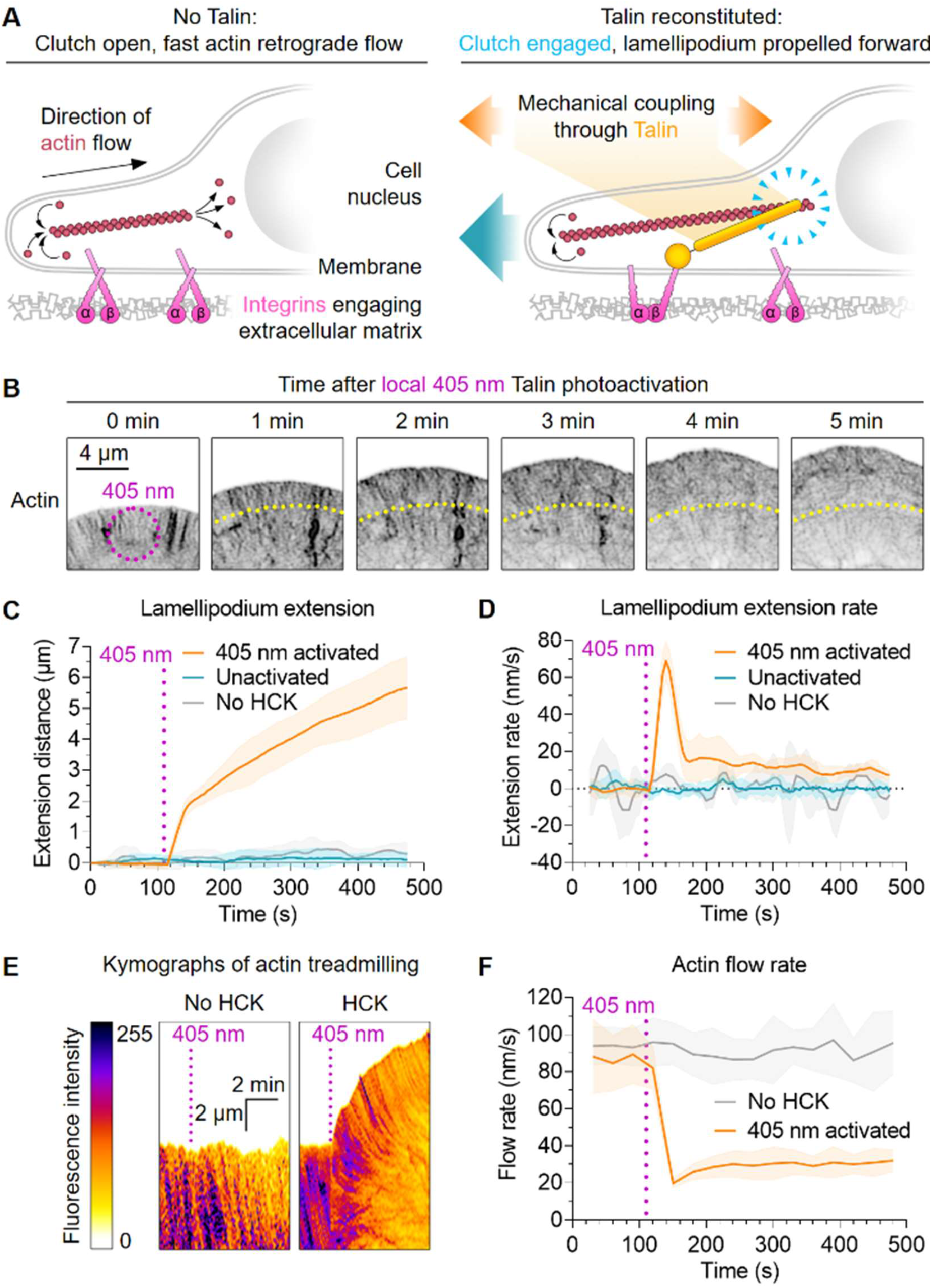
Photocontrol of SpyTag003/SpyCatcher003 reactivity in living cells. (A) Schematic of talin’s role as an adhesion clutch. (B) Photoactivation of cell spreading. Talin knock-out cells transfected with caged split talin were activated by 405 nm for 5 s (magenta ring) and imaged at the indicated time-points. Inverted LifeAct-mNeonGreen signal for actin is shown. Yellow indicates the original lamellipodium edge. (C-D) Quantification of lamellipodium extension distance and extension rate. Cells were activated as in B and imaged for LifeAct-mNeonGreen. n=5–11 cells. (E) Actin dynamics after photoactivation. Cells were activated as in B and kymographs created for lamellipodium LifeAct-mNeonGreen. (F) Quantification of actin treadmilling. Magenta indicates the point of 405 nm activation. Mean ± 1 SD, n=6–8 cells. Line represents the mean, with shading ± 1 SD.

Having confirmed robust photouncaging in lysates, we investigated talin reconstitution in fibroblasts with knock-out of both endogenous talin genes.^21^ Talin knock-out cells transfected with LifeAct-mNeonGreen-IRES-Talin head-SpyTag003, SpyCatcher003(K31HCK)-Talin rod-mScarletI and HCK RS plasmids were cultured with HCK and imaged by confocal microscopy with lasers at 405 nm (photoactivation), 488 nm (mNeonGreen, a bright green fluorescent protein) and 561 nm (mScarletI, a bright red fluorescent protein). Unactivated cells could not spread or polarize, consistent with the lack of functional talin.^21,23^ Local photoactivation at 405 nm for 5 s led to lamellipodia extension within seconds (Fig. 2B,C, Movie S1), indicating rapid reconstitution of talin in cells. We did not observe spreading of unactivated cells imaged at 488 nm (Fig. 2C, Movie S1), so typical microscopy conditions did not cause unintended photoactivation. Similarly, we did not observe spreading upon 405 nm exposure of cells transfected as above but with the equivalent DMSO concentration in place of HCK (Fig. 2C).

Upon talin reconstitution, we observed biphasic extension of lamellipodia, with a fast initial phase (∼70 nm/s) followed by a slower phase (10-20 nm/s) (Fig. 2C,D). Actin polymerization at the cell periphery is the main driving force propelling the lamellipodium forward, so we investigated actin treadmilling by tracking LifeAct-mNeonGreen (Fig. 2E). Unactivated cells had fast initial actin rearward flow at ∼90 nm/s (Fig. 2F). Talin reconstitution led to a sharp drop to ∼20 nm/s, followed by gradual recovery to ∼30 nm/s (Fig. 2F). The sharp drop in actin retrograde flow coincides with the phase of fast lamellipodium extension, suggesting the integrin-talin-actin clutch is rapidly engaged upon talin photoactivation.

Force-sensing by talin generates cellular signalling anisotropy, regulating cell polarization.^23^ Given the covalent SpyTag003:SpyCatcher003 interaction, this photoactivation strategy should allow extended cell polarization experiments covering tens of minutes. Talin knock-out fibroblasts transfected with talin head-SpyTag003, SpyCatcher003(K31HCK)-Talin rod-mScarletI and HCK RS plasmids were cultured with HCK and photoactivated with wide-field 405 nm light for 1 min. Cells were fixed at selected time-points and analyzed for cell area and morphology. Activated cells showed fast initial spreading and reached maximal area ∼10 min after photoactivation (Fig. 3A,B). In contrast, cell polarization was triggered only when the maximal cell area was reached and continued to develop until the end of the experiment (Fig. 3C). As expected, cells without HCK did not react to photoactivation (Fig. 3C,D).

**Figure 3.**
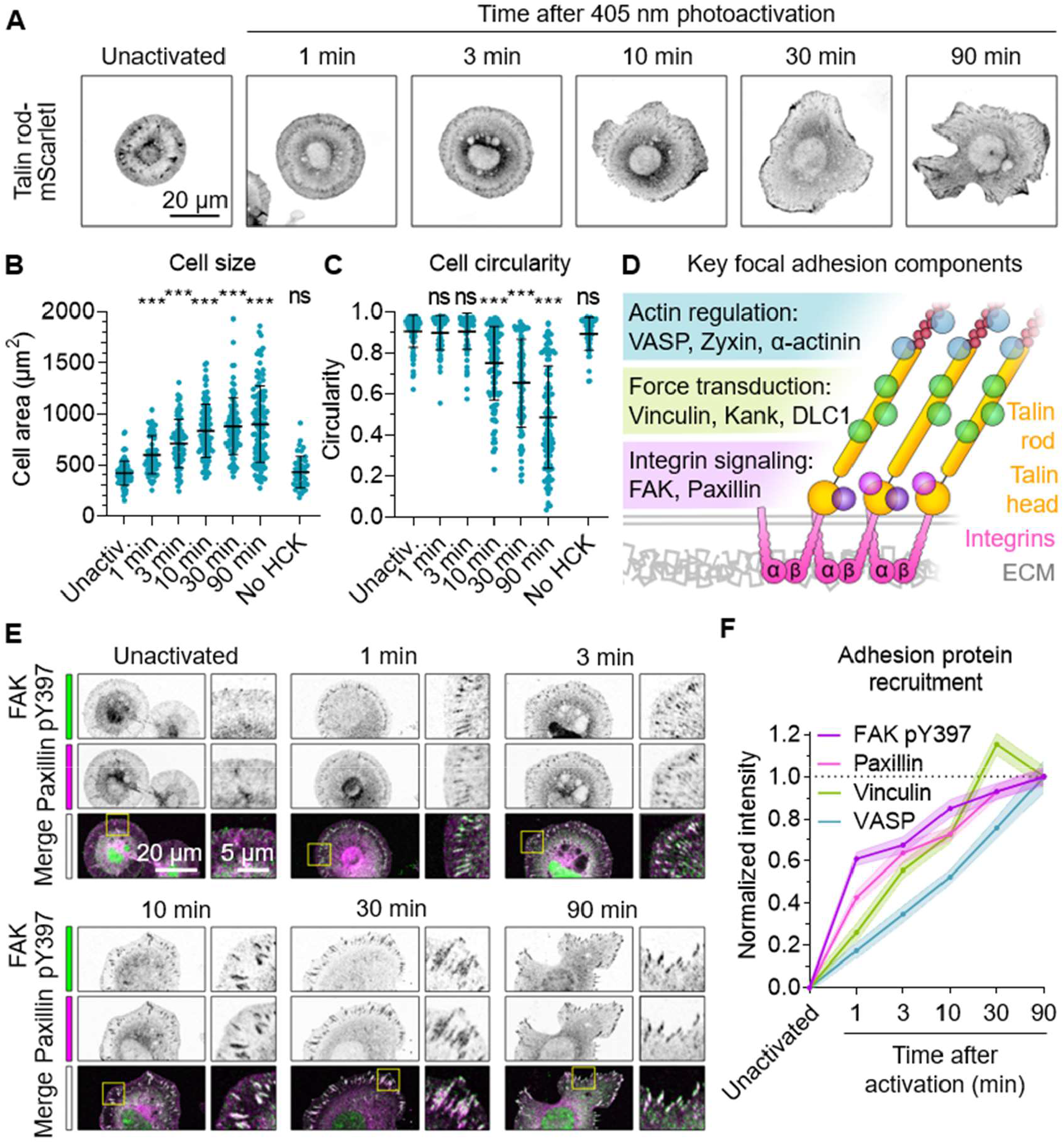
Photoactivation of talin allows precise control over adhesion complexes. (A) Cell spreading and polarization after photoactivation. Talin knock-out cells expressing talin head-SpyTag003, SpyCatcher003(K31HCK)-Talin rod-mScarletI and HCK RS were activated by 405 nm wide-field illumination for 1 min and fixed at the indicated time, before fluorescence microscopy for mScarletI. (B-C) Quantification of cell size and circularity following talin photoactivation as in A. Each blue circle is one cell, with black lines showing mean ± 1 SD. Compared with unactivated cells using One-way ANOVA and Dunnett’s test: *** p<0.0001, ns p>0.05, n=45–110 cells from 2 independent experiments. (D) Schematic of key adhesion components. Interactions with talin’s rod domain generate three functional layers (colored bars). (E) Recruitment of adhesion components after talin photoactivation. Talin knock-out cells were photoactivated as in A and stained with antibodies in fluorescence microscopy (zoom of the yellow square in the right image). Overlap of FAK pY397 (green) and paxillin (magenta) in the merge shows as white. (F) Quantification of recruitment to adhesions following talin photoactivation, as in E. Line represents the mean, with shading ± 1 SEM. n=60–85 adhesions in 12–17 cells from 2 independent experiments.

Stretching of talin rods regulates recruitment and release of many adhesion components, generating a structure with distinct functional layers (Fig. 3D).^28^ However, the heterogeneous and dynamic structure of adhesion complexes makes it challenging to define the temporal hierarchy of adhesion protein recruitment. Having validated our method for triggering synchronized adhesion, we investigated rates of recruitment for key adhesion components. We observed fast initial recruitment to adhesions of activated focal adhesion kinase (FAK) phosphorylated at tyrosine 397 (pY397), reaching half-maximal intensity <1 minute after photoactivation (Fig. 3E,F, Fig. S2B). Paxillin and vinculin reached half-maximal intensity at 3 minutes, with paxillin being slightly faster (Fig. 3E,F, Fig. S2A,C,D). In contrast, recruitment of the actin regulator vasodilator-stimulated phosphoprotein (VASP) reached half-maximal intensity only after 10 minutes (Fig. 3F, Fig. S2A,E).

## Conclusions

Robust optical control of protein complexation relies on sufficient bond life of the activated complex, ideally exceeding the natural turnover rate of the studied proteins. Interaction stability is especially challenging where the interface is under mechanical tension. Careful analysis of interface stability has allowed the use of elegant non-covalent optogenetic tools in reconstituting force-bearing proteins.^27,29^ However, local changes in force magnitude, duration and application rate can affect bond stability and lead to inconsistent or unrepresentative results. To overcome this limitation, we developed SpyCatcher003(K31HCK) for visible light photoactivation and demonstrated its application in the covalent reconstitution of split talin. Optical control of talin reconstitution allowed us to probe the time-scale of initial adhesion complex formation, revealing biphasic extension of lamellipodia upon engagement of the adhesion clutch. We also demonstrated the use of SpyCatcher003(K31HCK) coupling over a longer time-course, establishing a hierarchy of adhesion protein recruitment after engaging the adhesion clutch. The recruitment rates of adhesion proteins followed the layer structure of the adhesion complex (Fig. 3D),^28,30^ suggesting that talin not only governs the nanoscale organization of the adhesion but also the timing of protein recruitment.

Light control of covalent reactivity can also be achieved with bispecific molecules regulating the bridging of SNAP-tag (19 kDa) with HaloTag (33 kDa).^31,32^ However, the larger size of this protein pair may reduce the range of accessible sites. Split intein reaction may also be regulated by photocaged tyrosine but the reconstitution over 4 hours may limit applicability for cellular processes.^33^ While this work was in progress, a related approach was reported with photocaging of the slower first-generation SpyCatcher, using *o*-nitrobenzyloxycarbonyl-caged lysine.^34^ This approach used 365 nm wide-field uncaging with 20 min uncaging time. Hence, the 405 nm-responsive amino acid used here should have lower phototoxicity for cell biology studies.^17,35–37^ Also, 405 nm lasers are common on confocal microscopes, allowing uncaging at a spatiotemporal resolution not easily achieved using 365 nm wide-field light sources and photomasks.

Beyond adhesion, SpyCatcher003(K31HCK) may become a broadly applicable tool for photocontrol of biomolecules. A robust cellular response was initiated in seconds here, opening possibilities for spatiotemporal control of highly dynamic intracellular and extracellular processes.

## Acknowledgements

Funding was provided by Biotechnology and Biological Sciences Research Council (BBSRC BB/T004983/1, S.K.V. and M.H.) and Academy of Finland Postdoctoral researcher funding (339449, R.R.). We acknowledge Academy of Finland (331946, V.P.H.), Cancer Foundation Finland (V.P.H.), Sigrid Juselius Foundation (V.P.H.), and National Institutes of Health (R01AI175067, A.D.) for financial support. The authors acknowledge the Biocenter Finland and Tampere Imaging Facility for service and infrastructure support. We thank Prof. Reinhard Fässler (Max Planck Institute of Biochemistry) and Prof. Carsten Grashoff (University of Münster) for help with the talin knock-out fibroblasts.

## License information

For the purpose of Open Access, the author has applied a CC BY public copyright licence to any Author Accepted Manuscript (AAM) version arising from this submission.

## Supporting information

Supplementary Figure S1: Western blots for the validation of SpyCatcher003(K31HCK) photoactivation.

Supplementary Figure S2: Representative images of Vinculin and VASP recruitment, raw data for adhesion protein recruitment analysis.

Supplementary Movie S1: Time-lapse image series of cell spreading upon 405 nm photoactivation of talin reconstitution.

## TOC graphic

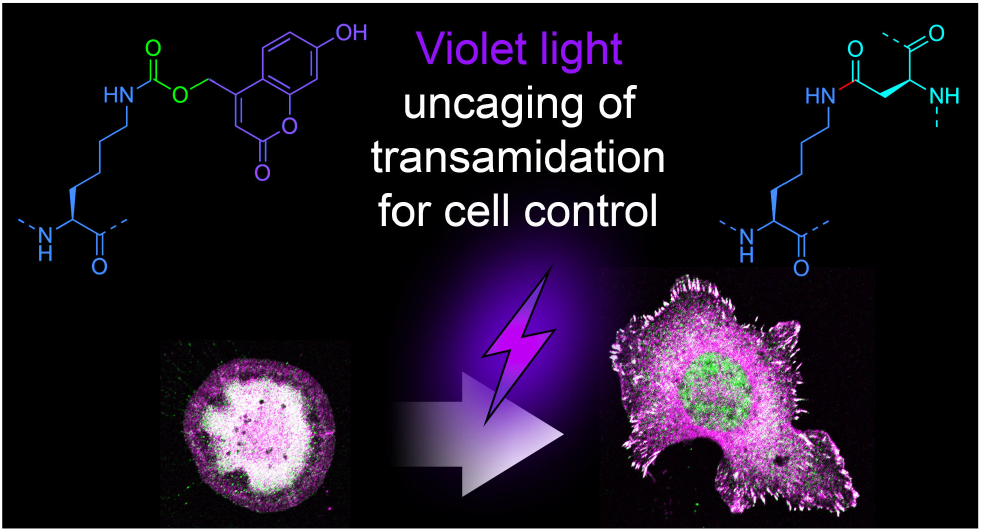

## Notes

### Competing Interest Statement

M.H. is an inventor on patents on spontaneous amide bond formation (EP2534484) and SpyTag003:SpyCatcher003 (UK Intellectual Property Office 1706430.4) and is a SpyBiotech co-founder and shareholder. All other authors have no competing interests to declare.

### Summary of Updates

Updated author list and abstract

