## Supplementary material for "Visible light-induced specific protein reaction delineates early stages of cell adhesion": Methods and Supp Figures

##### Plasmids and cloning

We used standard PCR methods with Q5 High-Fidelity 2× Master Mix (New England Biolabs) and Gibson assembly to perform cloning and site-directed mutagenesis. All open-reading frames were validated by Sanger sequencing (Source Bioscience). Residue numbers for SpyCatcher003 variants follow PDB ID: 2X5P.[38] Genscript plasmid cloning service was used to synthesize talin expression constructs for experiments in talin knock-out cells.

pcDNA3-mCherry-TAG-EGFP with one copy of *Desulfitobacterium hafniense* pyrrolysyl-tRNA$_{CUA}$ (PylT) with G8U under the control of a U6 promoter (1× U6-PylT) and one copy of *Methanosarcina barkeri* PylT with U25C under the control of a H1 promoter (1× H1-PylT) has been described.[39] *D. hafniense* PylT G8U mutation restores canonical U8-A14 interaction and *M. barkeri* PylT U25C mutation stabilizes the anticodon stem and slightly increases amino acid incorporation efficiency.[40,41] pcDNA3-EGFP-Talin head-SpyCatcher003 and pcDNA3-EGFP-Talin head-SpyCatcher003(K31TAG) (GenBank and Addgene deposition in progress) with 1× U6-PylT and 1× H1-PylT were derived from pcDNA3-mCherry-TAG-EGFP and pEGFP-C1-EGFP-Talin head-SpyCatcher003 (GenBank MN527523, Addgene ID 133566),[42] which have been described. Talin head comprises amino acids 1-433 of mouse talin-1. The amber stop codon TAG instead of the lysine at position 31 of SpyCatcher003 allows for the incorporation of 7-hydroxycoumarin-caged lysine (HCK) when co-expressed with HCK tRNA synthetase (HCK RS) and PylT.

pcDNA3-TfR-SpyCatcher003-superfolder green fluorescent protein (sfGFP) and pcDNA3-TfR-SpyCatcher003(K31TAG)-sfGFP (GenBank and Addgene deposition in progress) with 1× U6-PylT and 1× H1-PylT were derived from pcDNA3-mCherry-TAG-EGFP and pENTR4-TfR-sfGFP-SpyCatcher003 (GenBank MN433890, Addgene ID 133451),[42] which have been described. TfR comprises the Transferrin receptor transmembrane domain and cytosolic region with Y20C and F23A mutations to block internalization and allows for mammalian cell surface expression of SpyCatcher003.[42] In this context, sfGFP should only be expressed when HCK has successfully been incorporated in the growing polypeptide chain.

pcDNA3-LifeAct-mNeonGreen-IRES-Talin head-SpyTag003 (GenBank and Addgene deposition in progress) with 1× U6-PylT and 1× H1-PylT was derived from pcDNA3-EGFP-Talin head-SpyCatcher003 and encodes mouse codon-optimized LifeAct-mNeonGreen, followed by encephalomyocarditis virus (EMCV) internal ribosome entry site (IRES) and Talin head-SpyTag003. pcDNA3-Talin head-SpyTag003 was derived from pcDNA3-LifeAct-mNeonGreen-IRES-Talin head-SpyTag003 by removing the cassette encoding LifeAct-mNeonGreen and IRES. pEGFP-C1 SpyCatcher003(K31TAG)-Talin rod (434-2541)-mScarletI (GenBank and Addgene deposition in progress) was derived from pEGFP-C1 SpyTag003-Talin rod (434-2541)-mCherry (GenBank Accession No. MN527524 and Addgene plasmid ID 133567) previously described.[42] In this context, mScarletI should only be expressed when HCK has successfully been incorporated in the growing polypeptide chain.

pE323-HCK RS contains the engineered HCK tRNA synthetase with five copies of PylT from *Methanosarcina mazei* with U25C under the control of a U6 promoter (5× U6-PylT) and has been described.[43] pET28a-SpyTag003-MBP (GenBank MN433888, Addgene ID 133450) has been previously described.[42]

##### Hydroxycoumarin-caged lysine

HCK was synthesized as previously described[44] and made up at 100 mM in DMSO. HCK stock was stored at -20 °C protected from light and directly added into cell culture medium after transfection. For all No-HCK control samples, an equal volume of DMSO was added. Handling of HCK was performed under low light conditions in a tissue culture hood (with the hood light off and the room lights dimmed). During assays, HCK-incorporated samples were handled at ambient light but protected with aluminum foil and/or placed in a closed polystyrene ice box during incubations and waiting times.

To avoid possible phototoxicity of reactive oxygen species in the 405 nm activation of HCK in living cells, 0.25 mM trolox (Sigma-Aldrich) was added as an antioxidant.[45] 100 mM stock of trolox was prepared in 99.7 % (v/v) ethanol and stored at 4 °C.

**Expression of caged SpyCatcher003 in HEK293T cells**

HEK293T cells were maintained in complete Dulbecco's Modified Eagle Medium [DMEM (Sigma-Aldrich) supplemented with 10% (v/v) fetal bovine serum (FBS, Sigma-Aldrich), as well as 50 U/mL penicillin and 50 µg/mL streptomycin (Thermo Fisher)] under humidified conditions at 37 °C and 5% (v/v) $CO_2$. The day before transfection, 250,000 HEK293T cells were seeded in 2 mL complete DMEM in 6-well plates (Greiner). Cells were transfected with 2 µg plasmid DNA (1 part pcDNA3-U6H1-EGFP-Talin head-SpyCatcher003 WT/K31TAG and 2 parts pE323-HCK RS) using 2 µL jetOPTIMUS (Polyplus) according to the manufacturer's instructions. HCK was added to a final concentration of 0.25 mM after transfection. For control wells, 0.25% (v/v) DMSO was added. Cells were grown for 48 h, before Western blot analysis with the plates wrapped in aluminum foil.

**HEK293T cell lysis**

HEK293T cells in 6-well plates were washed three times with 1 mL phosphate-buffered saline (PBS: 137 mM NaCl, 2.7 mM KCl, 10 mM $Na_2HPO_4$, 1.8 mM $KH_2PO_4$) pH 7.4, then incubated in 200 µL ice-cold lysis buffer [25 mM Tris-HCl pH 7.5 (pH adjusted at 25 °C) + 150 mM NaCl + 5% (v/v) glycerol + 1% (v/v) Triton X-100, supplemented with cOmplete Mini EDTA-free Protease Inhibitor Cocktail (Roche) and 1 mM phenylmethylsulfonylfluoride (PMSF, Thermo Fisher)] for 10 min on ice in a closed ice box. The cells were scraped off, transferred to 1.5 mL tubes and centrifuged at 20,000 g for 10 min at 4 °C. The supernatant was used immediately or stored at -20 °C. Protein concentrations were determined by bicinchoninic acid assay (BCA Protein Assay Kit, Thermo Fisher) in 96-well format using 5 µL cell lysate diluted with 5 µL lysis buffer and correcting for lysis buffer, with bovine serum albumin (BSA) standards at 0, 125, 250, 500, 750, 1000, 1,500 or 2,000 µg/mL. Incubation with 200 µL BCA working solution was performed at 37 °C for 30 min. $A_{562}$ measurement was performed using a FLUOstar Omega plate reader with FLUOstar Omega software version 5.10 R2 (both BMG Labtech) at 25 °C.

**Initial optimization of caged SpyCatcher003 expression in HEK293T cells**

Optimization was performed in 24-well plates (Greiner). HEK293T cells were seeded at 50,000 cells/mL in 0.5 mL complete DMEM. Cells were transfected with 0.25 µg total DNA using 0.25 µL jetOPTIMUS, at molar ratios 1:1, 1:2 and 2:1 [pcDNA3-U6H1-TfR-SpyCatcher003(K31TAG)-sfGFP to pE323-HCK RS]. Directly after transfection, HCK was added to a final concentration of 0.25 mM, 0.5 mM or 1 mM. Western blot analysis was performed 48 h after transfection. For lysis, HEK293T cells were washed three times with 0.5 mL PBS pH 7.4, then incubated in 75 µL ice-cold lysis buffer and processed as described above.

**Protein purification by Ni-NTA**

SpyTag003-MBP was expressed in *E. coli* EXPRESS BL21(DE3) (Lucigen) and purified by Ni-NTA at 4 °C. Bacterial cell pellets of SpyTag003-MBP were thawed and resuspended in Ni-NTA buffer (50 mM Tris-HCl pH 7.8 + 300 mM NaCl) supplemented with cOmplete Mini EDTA-free Protease Inhibitor Cocktail (Roche) and 1 mM PMSF. Cells were lysed by sonication on ice at 50% duty cycle for 4 × 1 min with 1 min rest between runs. Cell lysates were clarified by centrifugation at 30,000 g for 30 min at 4 °C, then incubated with Ni-NTA agarose (Qiagen) for 1 h at 4 °C with rolling. Resin was pelleted by centrifugation, then washed with 20 column volumes (CV) of Ni-NTA buffer and Ni-NTA wash buffer (50 mM Tris-HCl pH 7.8 + 300 mM NaCl + 10 mM imidazole). The resin was transferred to Econo-Pac Chromatography Columns (Bio-Rad) and washed with a further 20 CV of Ni-NTA wash buffer by gravity flow. Elution was performed with Ni-NTA elution buffer (50 mM Tris-HCl pH 7.8 + 300 mM NaCl + 200 mM imidazole) by incubating the resin in 4 × 1 CV for 5 min. SpyTag003-MBP was dialyzed into PBS pH 7.4 in three dialysis steps using 3.5 kDa molecular weight cut-off (MWCO) dialysis tubing (Spectrum Labs). Protein concentrations were determined by $A_{280}$ measurement on a NanoDrop One (Thermo Fisher) using NanoDrop One software version 1.4.2, with extinction coefficients predicted by ExPASy ProtParam.[46]

**Uncaging and reaction of SpyCatcher003**

Equal amounts of protein lysates as determined by BCA assay were transferred to PCR tubes. To test spontaneous uncaging by ambient light, samples were incubated exposed to room light in a cold room or next to a window on ice for the indicated amounts of time. For uncaging, designated samples were irradiated on a ChemiDoc XRS+ using the Ethidium Bromide setting for the indicated amounts of time at 25 °C. Samples were transferred back to ice to chill and incubated with 50 μM SpyTag003-MBP at 4 °C for 1 h. Reaction was stopped by mixing with 6× SDS loading buffer [234 mM Tris-HCl pH 6.8, 24% (v/v) glycerol, 120 μM bromophenol blue, 234 mM SDS, supplemented with 60 mM dithiothreitol (DTT) for reduced samples] and heating at 95 °C for 5 min in a Bio-Rad C1000 thermal cycler.

**Western blot**

SDS-PAGE was performed using an XCell SureLock system (Thermo Fisher) with 10% (w/v) polyacrylamide gels. Gels were run in 24 mM Tris base, 192 mM glycine, 3.5 mM SDS at 200 V. Proteins were transferred onto nitrocellulose membranes by dry transfer using an iBlot 2 gel transfer device (Thermo Fisher) at 25 V for 10 min. Membranes were blocked for 1 h at 25 °C in 5% (w/v) skimmed milk in PBS. The primary antibody was diluted in 2.5% (w/v) skimmed milk in 0.05% PBST [PBS pH 7.4 + 0.05% (v/v) Tween 20], with mouse anti-GFP (Thermo Fisher, MA5-15256, clone GF28R) at 1:2,000 or mouse anti-SpyCatcher003 serum[47] at 1:300. Incubation was performed overnight at 4 °C. Membranes were washed three times in 0.1% PBST [PBS pH 7.4 + 0.1% (v/v) Tween 20] for 5 min at 25 °C. Secondary antibody detection was performed by staining with goat anti-mouse IgG Horseradish Peroxidase (Sigma-Aldrich, A4416) at 1:5,000 for 1 h at 25 °C in 2.5% (w/v) skimmed milk in 0.05% PBST. After three further washes in 0.1% PBST and one wash in PBS at 25 °C, membranes were developed at 25 °C with SuperSignal West Pico Plus Chemiluminescent Substrate (Thermo Fisher) and imaged using a ChemiDoc XRS+ imager (Bio-Rad). In immunoblotting with anti-EGFP antibody, we observed EGFP-containing degradation products of EGFP-Talin head-SpyCatcher003(K31TAG), consistent with reported high stability of EGFP in cells.[48] A colon is used to indicate products linked by Tag/Catcher isopeptide bond formation.

**Expression of caged SpyCatcher003 in talin knock-out cells**

*Tln1-/-Tln2-/-* mouse kidney fibroblasts were a kind gift from Prof. Carsten Grashoff, University of Münster, and have been previously described.[49] Talin knock-out cells were maintained in complete FluoroBrite medium [FluoroBrite DMEM (A1896701, Thermo Fisher) supplemented with 10% (v/v) FBS (10270106, Gibco) and 1% (v/v) GlutaMax (35050061, Thermo Fisher)] under humidified conditions at 37 °C and 5% (v/v) $CO_2$. The cell line was regularly tested for mycoplasma contamination.

Talin knock-out cells were transiently transfected with Neon Transfection System electroporator (Thermo Fisher) according to the manufacturer's instructions. Briefly, $10^6$ cells in 100 µL were transfected with 12.5 µg of total plasmid DNA at 2:2:1 molar ratio of pE323-HCK RS, SpyCatcher003(K31TAG)-Talin rod (434-2541)-mScarletI and Talin head-SpyTag003 plasmids, respectively. The electroporation parameters were 1,400 V, 30 ms and 1 pulse.

**Live-cell imaging and on-stage 405 nm photoactivation**

Zeiss high-performance 170-µm-thick coverslips were washed with 2% (v/v) Hellmanex-III (Sigma-Aldrich) in a bath sonicator at 40 °C for 20 min (Finnsonic), rinsed with deionized water and sonicated again in 96% (v/v) ethanol. Coverslips were rinsed with deionized water, air-dried, and attached to perforated 35 mm polystyrene dishes (MatTek). Attached coverslips were coated with 25 µg/mL human fibronectin in PBS (pH 7.4) for 30 min at 37 °C and washed twice with PBS (pH 7.4). Fibronectin was purified from human plasma preparate (Octaplas) using gelatin affinity chromatography (Gelatin Sepharose 4B; GE Healthcare) as previously described.[50]

Transfected talin knock-out cells were plated on glass bottom dishes (166,000 cells per dish) in 120 µL of complete FluoroBrite medium and incubated at 37 °C and 5% (v/v) $CO_2$ for 1 h to allow cell attachment. Dishes were filled with complete FluoroBrite media and the incubation at 37 °C and 5% (v/v) $CO_2$ continued for 2 h. Media was replaced with 800 µL of FluoroBrite-trolox-HCK (Complete FluoroBrite supplemented with 0.25 mM trolox and 0.25 mM HCK) and cells incubated at 37 °C and 5% (v/v) $CO_2$ for 16-18 h. Before imaging, media was replaced with FluoroBrite-trolox (Complete FluoroBrite media supplemented with 0.25 mM trolox) and the dish mounted to a humified 37 °C and 5% (v/v) $CO_2$ incubator on the microscope stage.

For live-cell imaging and on-stage 405 nm photoactivation, we used a Nikon Eclipse Ti2-E inverted microscope equipped with A1R+ laser scanning confocal (Nikon), Perfect Focus System (Nikon) and Nikon SR HP Plan Apo 100×/1.35 Silicone immersion objective. Single cells were imaged at 80 nm pixel size using 488 nm solid state laser (0.25 - 0.5% laser) for mNeonGreen excitation and 561 nm solid state laser (0.2 - 0.5% laser) for mScarletI excitation. Differential Interference Contrast (DIC) images were captured using transmitted 488 nm excitation light. Images were captured at 5 s intervals for 23 images before photoactivation. For HCK photoactivation, a circular region of 4.0 µm was activated for 5 loops (4.8 s total) with 10.5 µW (5%) 405 nm laser. Imaging was then immediately continued for 73 images at 5 s intervals.

"Unactivated" cells were cultured with HCK but not 405 nm activated. In Fig. 3, "No HCK" cells were cultured without HCK, activated with 405 nm, and allowed to spread for 90 min. When green and magenta images are merged, the overlapping regions appear white.

**Lamellipodium extension analysis**

Fiji distribution of ImageJ 1.53t[51] was used to analyze lamellipodium extension after HCK photoactivation. A 10-pixel (0.8 µm) wide line perpendicular to the cell edge was drawn across the cell lamellipodium at the site of photoactivation. A kymograph of mNeonGreen intensity was created along this line, covering all 96 frames of the time-series. Using free-hand selection tool, the movement of the lamellipodium edge over time was traced in the kymograph by following the shift in the position of mNeonGreen intensity. The selection created in this way was used to quantify the number of pixels above the line (inside cell) and below the line (outside cell) for all columns of pixels, each representing a single image captured at 5 s intervals. For relative lamellipodium extension, the results were normalized with the initial position of the lamellipodium edge. To calculate the lamellipodium extension rate, sharp irregularities in the extension data were smoothed by averaging with a 4-frame sliding window and the rate of change over time was calculated.

**Actin flow-rate analysis**

To improve the clarity of LifeAct-mNeonGreen live-cell imaging data, Huygens Essential 23.04 (Scientific Volume Imaging) with classic maximal likelihood estimation (CMLE) deconvolution algorithm was used. Deconvoluted images were analyzed using Fiji distribution of ImageJ 1.53t.[51] Briefly, at the site of photoactivation, a 20-pixel (1.6 µm) wide line perpendicular to the cell edge was drawn across the lamellipodium to create 5 parallel kymographs, each representing the mean intensity of 4 pixels in the original image. These kymographs were used to measure manually the slope of actin flow at 30 s (6 pixel) intervals using the line selection tool. Of the 5 parallel kymographs, at each time point, the one with the best image clarity was selected for analysis. The slopes measured in this way were used to calculate actin flow rate (nm/s) using trigonometry.

**Widefield 405 nm activation of talin knock-out cells**

Transfected talin knock-out cells were plated on 60 mm dishes (Nunc) in complete FluoroBrite and allowed to recover for 3 h at 37 °C and 5% (v/v) $CO_2$. The media was replaced with FluoroBrite-trolox-HCK and the incubation at 37 °C and 5% (v/v) $CO_2$ was continued for 16-18 h. After the addition of FluoroBrite-trolox-HCK, all subsequent culture steps were performed in low light conditions. Cells were rinsed with PBS (pH 7.4), detached using TrypLE Express (Thermo Fisher) and plated in FluoroBrite-trolox-HCK on fibronectin coated glass-bottom dishes, prepared as described above. After 1 h incubation at 37 °C and 5% (v/v) $CO_2$, cells were rinsed once with FluoroBrite-trolox and incubated at 37 °C and 5% (v/v) $CO_2$ for 15 min to wash out residual HCK. For HCK uncaging, dishes were placed on top of 2 × 6 W 405 nm LED panels (Sovol) with 45 mW/cm$^2$ total intensity and photoactivated for 60 s. Photoactivated cells were incubated at 37 °C and 5% (v/v) $CO_2$ and fixed at defined timepoints. For fixation, media was aspirated and 4% (w/v) paraformaldehyde in PBS (pH 7.4) immediately added for 20 min at 22 °C. Cells were washed 4 times with PBS (pH 7.4) and stored at 4 °C.

**Immunostaining**

Fixed cells were permeabilized with 0.2% (v/v) Triton X-100 in PBS (pH 7.4) for 5 min at 22 °C. Non-specific antibody binding was inhibited with blocking buffer [5% (v/v) fetal calf serum (Gibco), 1% (w/v) bovine serum albumin (Sigma-Aldrich) and 0.05% (v/v) Triton X-100 (Sigma-Aldrich) in PBS (pH 7.4)] for 30 min at 22 °C. Primary antibodies were diluted in the blocking buffer and incubated for 90 min at 22 °C at the following dilutions: anti-vinculin V9131 (Sigma-Aldrich, RRID:AB_477629) 1:400; anti-VASP HPA005724 (Sigma-Aldrich, RRID:AB_1858721) 1:100; anti-FAK pY397 ab81298 (Abcam, RRID:AB_1640500) 1:200; anti-paxillin BD610051 (BD Biosciences, RRID:AB_397463) 1:100. Cells were washed 4 times for 10 min with PBS (pH 7.4). Fluorescently-labelled secondary antibodies were diluted in the blocking buffer at 1:250 and incubated on samples for 90 min: Goat anti-Mouse IgG (H+L) AlexaFluor Plus 488 A32723 (Thermo Fisher, RRID:AB_2633275), Goat anti-Rabbit IgG (H+L) AlexaFluor Plus 488 A32731 (Thermo Fisher, RRID:AB_2633280), Goat anti-Mouse IgG (H+L) AlexaFluor Plus 647 A32728 (Thermo Fisher, RRID:AB_2633277), Goat anti-Rabbit IgG (H+L) AlexaFluor Plus 647 A32733 (Thermo Fisher RRID:AB_2633282).

Cells were washed 4 times for 10 min with PBS (pH 7.4) and covered with a clean coverslip using Prolong Glass antifade mounting media (Thermo Fisher). The mounting media was cured at 24 °C for a minimum of 40 h before imaging.

**Cell size and morphology analysis**

For cell size and morphology analysis, mScarletI signal of fixed samples was imaged using a Nikon Eclipse Ti2-E inverted microscope equipped with A1R+ laser scanning confocal unit (Nikon) and Nikon Apo 60×/1.40 Oil λS DIC N2 objective. For *No HCK* samples (Fig. 3B,C) not expressing mScarletI because of polypeptide chain termination, anti-vinculin antibody staining was imaged instead.

Cell area and circularity were analyzed in Fiji distribution of ImageJ 1.53t[51] using the free-hand selection tool to trace cell boundaries. Cell circularity was calculated with the formula: circularity = $4\pi(\text{area}/\text{perimeter}^2)$. Cells with partially detached lamellipodium or extremely high mScarletI signal were excluded from the analysis. For each timepoint, 45 to 110 cells from 2 independent experiments were analyzed.

**Adhesion intensity analysis**

For adhesion intensity analyses, immunostained cells were imaged using a Zeiss Axio Observer.Z1 equipped with Zeiss LSM800 laser scanning confocal unit and Zeiss Plan-Apochromat 63×/1.40 oil immersion objective. 107 µm × 107 µm fields were imaged at 70 nm pixel size and 200 nm Z-stack interval, maintaining fixed laser intensities and detector gains for all samples.

Adhesion protein localization to adhesion sites was analyzed using Fiji distribution of ImageJ 1.53t.[51] The Z-stack slices with the highest adhesion/cytosol contrast were selected for the analysis. For each cell, the 5 adhesions with the highest fluorescence intensity were masked using freehand selection tool and the mean intensity of each selected region measured. For each image, the intensity of background control region outside the cell was measured and its intensity value was subtracted from all adhesion intensity values of the same image. For Figure 3F, adhesion intensity values were normalized by the mean intensity measured at the 90 min timepoint. For unactivated cells, no clear adhesion structures were observed and the reported intensity values represent mean lamellipodium intensity. For each adhesion protein, 60-85 adhesions in 12-17 cells from two independent experiments were analyzed. Statistical significance of results was tested using one-way ANOVA with Dunnett's post-test in GraphPad Prism 9.0.0 (GraphPad Software). All other samples were compared with the unactivated samples. p-values below 0.05 were considered statistically significant. p-values above 0.05 were considered non-significant (ns). Samples in the same experiment were prepared, imaged and analyzed under uniform conditions.

**Data availability**

Sequences of constructs are available in GenBank, as described in the section "Plasmids and Cloning". Indicated plasmids will be deposited before publication in the Addgene repository (https://www.addgene.org/Mark_Howarth/). Further information and request for resources and reagents should be directed to and will be fulfilled by the lead contact, M.H.. Requests for unnatural amino acid should be directed to A.D..

#### References

(38) Oke, M.; Carter, L. G.; Johnson, K. A.; Liu, H.; McMahon, S. A.; Yan, X.; Kerou, M.; Weikart, N. D.; Kadi, N.; Sheikh, M. A.; Schmelz, S.; Dorward, M.; Zawadzki, M.; Cozens,

C.; Falconer, H.; Powers, H.; Overton, I. M.; Van Niekerk, C. A. J.; Peng, X.; Patel, P.; Garrett, R. A.; Prangishvili, D.; Botting, C. H.; Coote, P. J.; Dryden, D. T. F.; Barton, G. J.; Schwarz-Linek, U.; Challis, G. L.; Taylor, G. L.; White, M. F.; Naismith, J. H. The Scottish Structural Proteomics Facility: Targets, Methods and Outputs. *J. Struct. Funct. Genomics,* **2010**, *11* (2), 167–180. https://doi.org/10.1007/s10969-010-9090-y.
(39) Zhou, W.; Wesalo, J. S.; Liu, J.; Deiters, A. Genetic Code Expansion in Mammalian Cells: A Plasmid System Comparison. *Bioorg. Med. Chem,* **2020**, *28* (24), 115772. https://doi.org/10.1016/j.bmc.2020.115772.
(40) Schmied, W. H.; Elsässer, S. J.; Uttamapinant, C.; Chin, J. W. Efficient Multisite Unnatural Amino Acid Incorporation in Mammalian Cells via Optimized Pyrrolysyl tRNA Synthetase/tRNA Expression and Engineered ERF1. *J. Am. Chem. Soc,* **2014**, *136* (44), 15577–15583. https://doi.org/10.1021/ja5069728.
(41) Herring, S.; Ambrogelly, A.; Polycarpo, C. R.; Söll, D. Recognition of Pyrrolysine tRNA by the Desulfitobacterium hafniense Pyrrolysyl-tRNA Synthetase. *Nucleic Acids Res*, **2007**, *35* (4), 1270–1278. https://doi.org/10.1093/nar/gkl1151.
(42) Keeble, A. H.; Turkki, P.; Stokes, S.; Anuar, I. N. A. K.; Rahikainen, R.; Hytönen, V. P.; Howarth, M. Approaching Infinite Affinity through Engineering of Peptide-Protein Interaction. *Proc. Natl. Acad. Sci. U.S.A,* **2019**, *116* (52), 26523–26533. https://doi.org/10.1073/pnas.1909653116.
(43) Zhou, W.; Hankinson, C. P.; Deiters, A. Optical Control of Cellular ATP Levels with a Photocaged Adenylate Kinase. *ChemBioChem,* **2020**, *21* (13), 1832–1836. https://doi.org/10.1002/cbic.201900757.
(44) Luo, J.; Uprety, R.; Naro, Y.; Chou, C.; Nguyen, D. P.; Chin, J. W.; Deiters, A. Genetically Encoded Optochemical Probes for Simultaneous Fluorescence Reporting and Light Activation of Protein Function with Two-Photon Excitation. *J. Am. Chem. Soc,* **2014**, *136* (44), 15551–15558. https://doi.org/10.1021/ja5055862.
(45) Ramakrishnan, P.; Maclean, M.; MacGregor, S. J.; Anderson, J. G.; Grant, M. H. Cytotoxic Responses to 405 nm Light Exposure in Mammalian and Bacterial Cells: Involvement of Reactive Oxygen Species. *Toxicol. in Vitro,* **2016**, *33,* 54–62. https://doi.org/10.1016/j.tiv.2016.02.011.
(46) Gasteiger, E.; Hoogland, C.; Gattiker, A.; Duvaud, S.; Wilkins, M. R.; Appel, R. D.; Bairoch, A. Protein Identification and Analysis Tools on the ExPASy Server. In *The Proteomics Protocols Handbook*; Walker, J. M., Ed.; Humana Press Inc., Totowa, NJ, **2005**; pp 571–608.
(47) Rahikainen, R.; Rijal, P.; Tan, T. K.; Wu, H.-J.; Andersson, A.-M. C.; Barrett, J.; Bowden, T.; Draper, S.; Townsend, A. R.; Howarth, M. Overcoming Symmetry Mismatch in Vaccine Nanoassembly via Spontaneous Amidation. *Angew. Chem. Int. Ed,* **2020**, anie.202009663. https://doi.org/10.1002/anie.202009663.
(48) Reichard, E. L.; Chirico, G. G.; Dewey, W. J.; Nassif, N. D.; Bard, K. E.; Millas, N. E.; Kraut, D. A. Substrate Ubiquitination Controls the Unfolding Ability of the Proteasome. *J. Biol. Chem*, **2016**, *291* (35), 18547–18561. https://doi.org/10.1074/jbc.M116.720151.
(49) Theodosiou, M.; Widmaier, M.; Böttcher, R. T.; Rognoni, E.; Veelders, M.; Bharadwaj, M.; Lambacher, A.; Austen, K.; Müller, D. J.; Zent, R.; Fässler, R. Kindlin-2 Cooperates with Talin to Activate Integrins and Induces Cell Spreading by Directly Binding Paxillin. *eLife,* **2016**, *5,* e10130. https://doi.org/10.7554/eLife.10130.
(50) Ruoslahti, E.; Hayman, E. G.; Pierschbacher, M.; Engvall, E. Fibronectin: Purification, Immunochemical Properties, and Biological Activities. *Meth. Enzymol,* **1982**, *82 Pt A,* 803–831. https://doi.org/10.1016/0076-6879(82)82103-4.
(51) Schindelin, J.; Arganda-Carreras, I.; Frise, E.; Kaynig, V.; Longair, M.; Pietzsch, T.; Preibisch, S.; Rueden, C.; Saalfeld, S.; Schmid, B.; Tinevez, J.-Y.; White, D. J.;

Hartenstein, V.; Eliceiri, K.; Tomancak, P.; Cardona, A. Fiji: An Open-Source Platform for Biological-Image Analysis. *Nat. Methods*, **2012**, *9* (7), 676–682. https://doi.org/10.1038/nmeth.2019.

### Supporting Figures

#### Visible light-induced specific protein reaction delineates early stages of cell adhesion

Rolle Rahikainen[a], Susan K. Vester[b,c], Paula Turkki[a], Chasity P. Janosko[d], Alexander Deiters[d], Vesa P. Hytönen[a], and Mark Howarth[b,e,]

[a]Faculty of Medicine and Health Technology, Tampere University, Arvo Ylpön katu 34, 33520, Tampere, Finland and Fimlab Laboratories, Biokatu 4, 33520, Tampere, Finland.
[b]Department of Biochemistry, University of Oxford, South Parks Road, Oxford, OX1 3QU, UK.
[c]Current address: Randall Centre for Cell and Molecular Biophysics, King's College London, New Hunt's House, London, SE1 1UL, UK.
[d]Department of Chemistry, University of Pittsburgh, Pittsburgh, PA 15260, United States.
[e]Department of Pharmacology, University of Cambridge, Tennis Court Road, Cambridge, CB2 1PD, UK.

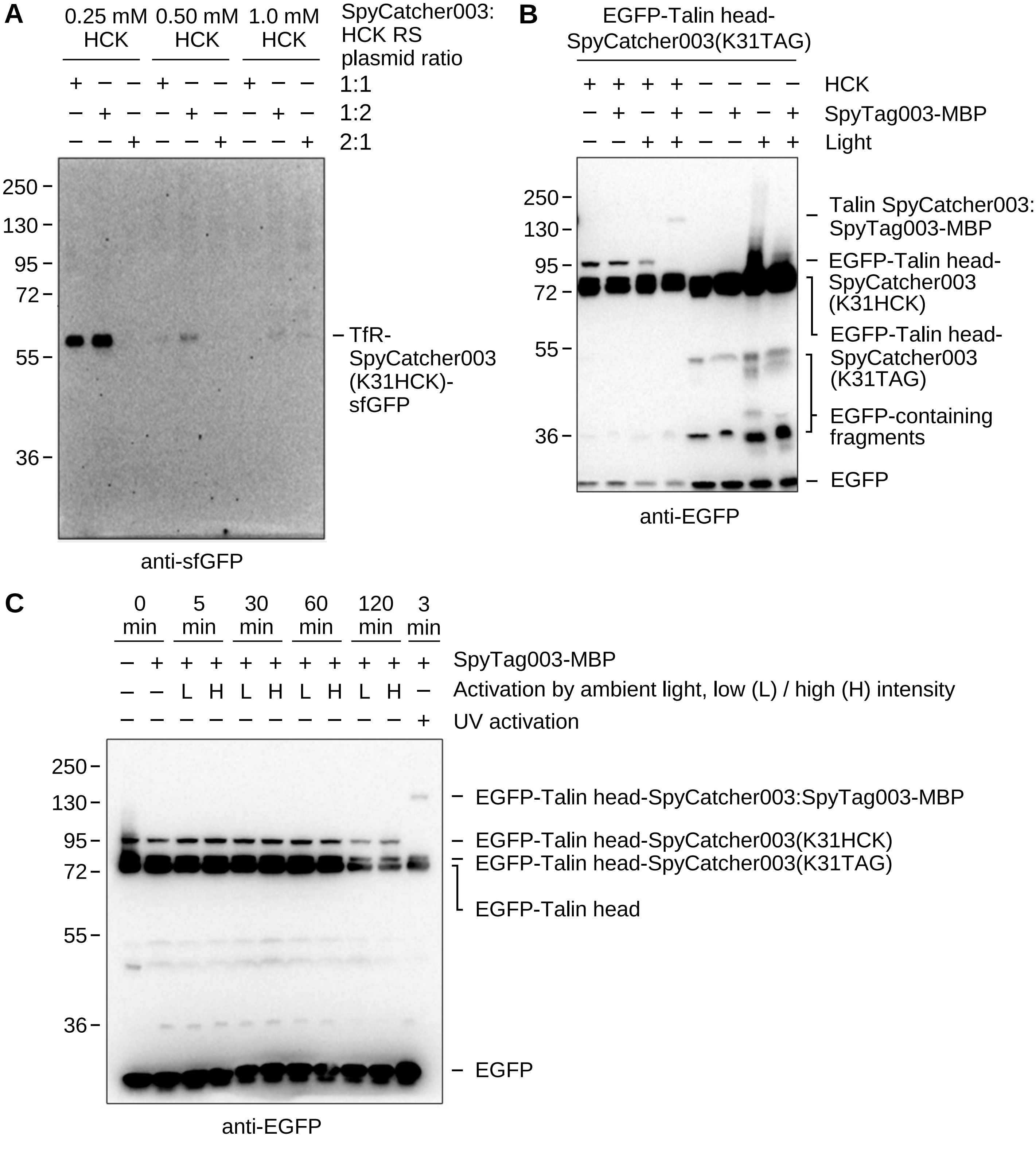


**Figure S1. Validation of SpyCatcher003(K31HCK) photoactivation.** (**A**) Titration of HCK concentration and plasmid ratios. HEK293T cells were transfected with different molar ratios of TfR-SpyCatcher003(K31TAG)-sfGFP and HCK RS plasmids and cultured in the presence of 0.25 mM – 1.0 mM HCK. Cell lysate was blotted for sfGFP, indicating successful incorporation of HCK in the polypeptide chain. GFP variants are highly stable to proteasomal degradation, so GFP-linked fragments are commonly observed to accumulate in cells.[48] (**B**) Photoactivation of covalent reactivity in EGFP-Talin head- SpyCatcher003(K31TAG), analyzed by Western blot against EGFP, following Fig. 1E. A colon is used to indicate isopeptide-linked products. (**C**) Photostability test for SpyCatcher003(K31HCK) in ambient light. Lysates of HEK293T cells were mixed with 50 µM SpyTag003-MBP and incubated for the indicated time, either in a dimly lit cold-room (Low intensity) or by the lab window (High intensity). The positive control sample was activated with a UV transilluminator.

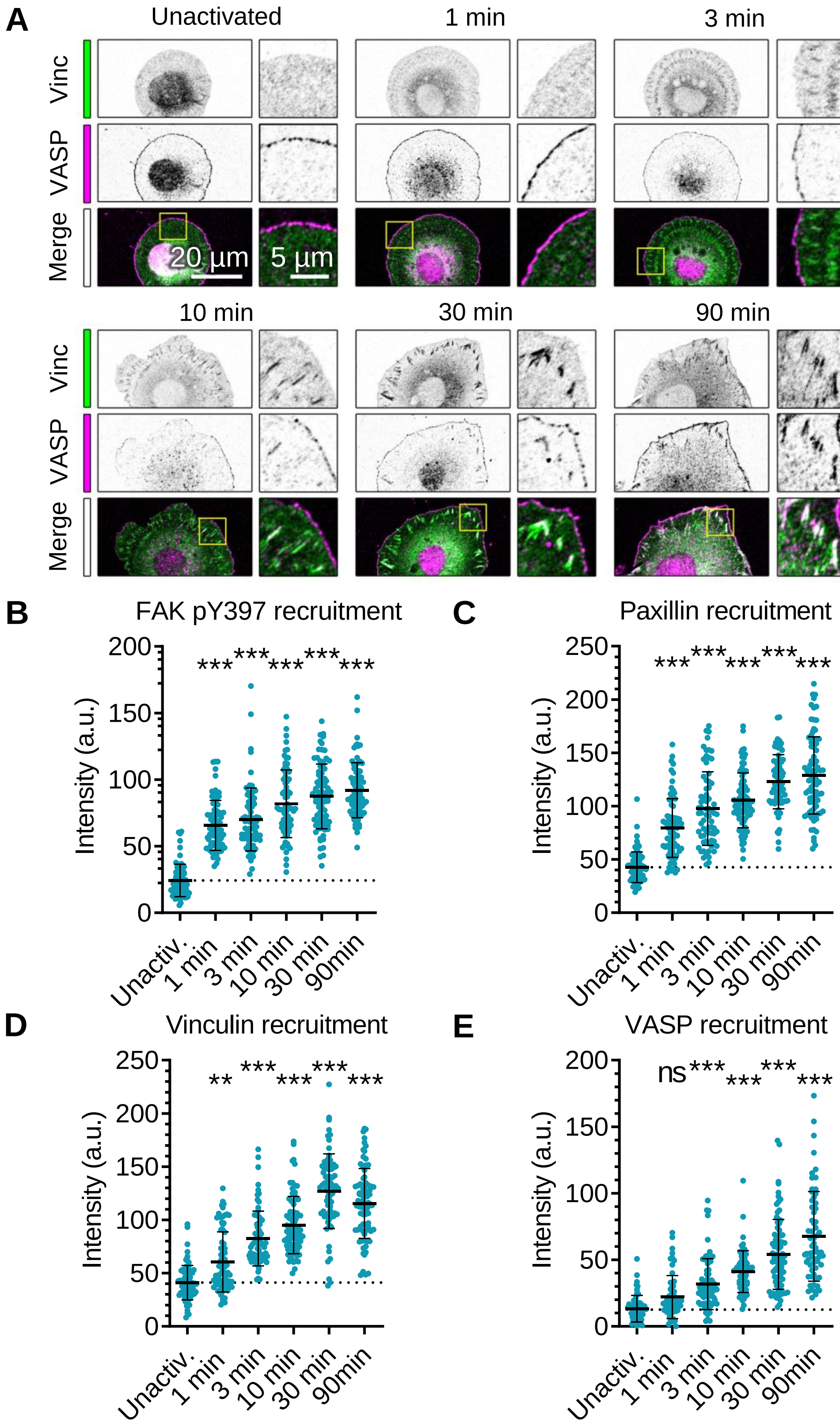


**Figure S2. Time-course for adhesion assembly.** (**A**) Vinculin and VASP recruitment after talin photoactivation. Talin knock-out cells were transfected as in Fig. 3E, activated at 405 nm and incubated for the indicated time. Cells were immunostained for VASP and vinculin. The yellow square is expanded in the right image. Vinculin and VASP are shown on their own in grayscale. Overlap of vinculin (green) and VASP (magenta) in the merge shows as white. (**B-E**) Recruitment of adhesion components following photoactivation. Raw adhesion intensities in arbitrary units (a.u.), to accompany normalized intensity in Fig. 3F. Each dot represents a single adhesion with mean ± 1 SD indicated as horizontal lines. FAK pY397: n = 75–81 adhesions in 15–16 cells. Paxillin: n = 70–86 adhesions in 14–17 cells. Vinculin: n = 74–81 adhesions in 12–15 cells. VASP: n = 60–86 adhesions in 12–15 cells. Results were pooled from two independent experiments. One-way ANOVA with Dunnett's test. *** $p < 0.0001$, ** $p < 0.001$, ns $p > 0.05$

**Movie S1 legend.**

Movie of split talin photoactivation in live cells. Talin knock-out cells were transfected with LifeAct-mNeonGreen, Talin head-SpyTag003, SpyCatcher003(K31HCK)-Talin rod-mScarletI and HCK RS and cultured with 0.25 mM HCK for 16–18 h. Left column shows inverted signal of actin labeling with LifeAct-mNeonGreen, middle column shows bright-field, and right column shows a merge of bright-field (grayscale) with LifeAct-mNeonGreen (green) determined by confocal microscopy. Top row: at 1 min 50 s, a circular 4-µm region was photoactivated by 405 nm illumination for 5 s, indicated by a magenta circle. Bottom row: cells treated in the same way but without 405 nm illumination. Photoactivation triggers initial local extension of the lamellipodium, followed by global cell spreading as reconstituted talin diffuses throughout the cell. The scale bar is 10 µm and the movie is shown at 6 frames per s (30x sped-up).
